# Natural killer cell TGF-β signaling regulates senolytic activity and vascular patterning in the postnatal lung

**DOI:** 10.1101/2025.08.29.673190

**Authors:** Declan J. Gainer, Kassandra M. Coyle, Matthew T. Rätsep, Douglas Quilty, Brian Tran, Sofia Skebo, M. Martin VandenBroek, Kimberly L. Laverty, Yupu Deng, Shawyon P. Shirazi, Hugh JM Brady, Jennifer M.S. Sucre, Eric Vivier, Niraj Shrestha, Hing C. Wong, Duncan J Stewart, Nicolle J. Dominik, Mark L. Ormiston

## Abstract

**Background:** Bronchopulmonary dysplasia (BPD) is a disease of neonatal lung development that is linked to impaired pulmonary vascularization, dysregulated transforming growth factor-β (TGF-β) signaling and the accumulation of senescent cells. Despite the established role for TGF-β signaling in promoting vascular remodeling and suppressing the senolytic activity of natural killer (NK) cells, the contribution of NK cell TGF-β signaling to postnatal lung patterning and the pathogenesis of BPD remains unclear.

**Methods:** Mice bearing an NK cell-selective deletion of the type-II TGF-β receptor (*Tgfbr2^NK-/-^*) were analyzed for vascular and alveolar structure, lung NK cell infiltration, senescence markers and lung function testing across neonatal and adult timepoints. Single-cell RNA sequencing of lung tissue from both neonatal mice and human infants with BPD was performed. The effect of enhanced NK cell activity in a hyperoxia-induced model of BPD was assessed in *Tgfbr2^NK-/-^*neonates, as well as pharmacologically, using the TGF-β ligand trap/IL-15 superagonist, HCW9218.

**Results:** Neonatal *Tgfbr2^NK-/-^* mice exhibited a baseline reduction in distal arteriolar density, impaired alveolarization, and sex-specific deficits in long-term lung function. Single-cell RNA sequencing identified the excessive clearance of senescent endothelial cells by TGF-β insensitive NK cells in the lungs of *Tgfbr2^NK-/-^* neonates, which served as a contributor of the BPD-like phenotype observed in naïve animals. *Tgfbr2^NK-/-^* mice were protected from impaired lung development in the hyperoxia model. Sequencing from lung tissue from infants with BPD confirmed excessive TGF-β signaling and cytotoxic impairment in NK cells. Treatment with HCW9218 prevented senescent cell accumulation and rescued lung development in the hyperoxia mouse model.

**Conclusions:** These findings identify TGF-β as a tunable regulator of NK cell senolytic activity that is essential to normal postnatal lung development. Excessive NK cell TGF-β signaling contributes to impaired lung development following exposure to neonatal hyperoxia and may serve as a viable therapeutic target for human BPD.

## Introduction

Vascular and airway development are highly integrated processes in the lung^1^. While primordial airways guide pulmonary vascular patterning in early embryonic lung development, these roles are reversed in the distal lung, where growth of the pulmonary arterioles and capillaries is critical to alveolarization^2^. This interdependency is best exemplified by bronchopulmonary dysplasia (BPD), a disease of impaired alveolar expansion that is strongly linked to the blunted development of the pulmonary circulation^3,4^. BPD can arise from various factors, including antenatal inflammation and postnatal hyperoxia secondary to pre-term birth^5^, and is a leading cause of morbidity and mortality in premature infants.

Dysregulated transforming growth factor-β (TGF-β) signaling has been identified as a contributor to BPD pathogenesis in both ventilatory and hyperoxic models of the disease^6–8^. Canonical TGF-β signaling involves phosphorylation of the transcriptional mediators, SMAD2 and SMAD3, which complex with SMAD4 and translocate to the nucleus to induce the expression of target genes involved in proliferation and differentiation^9^. Although TGF-β signaling is critical to proper lung patterning across various stages of development^10^, multiple studies have highlighted the pathological effects of excessive TGF-β on pulmonary stromal populations and associated connective tissue in BPD^11,12^. In contrast, the impact of TGF-β signaling on lung-resident immune populations, and the downstream consequences of this signaling on lung development, are less clearly defined.

In natural killer (NK) cells, TGF-β suppresses maturation, proliferation, cytotoxic function and, more recently, has been shown to promote the conversion of cytotoxic NK cells towards tissue-resident or ILC1-like phenotypes^13^. These specialized NK cell subsets are known to influence tissue homeostasis in a range of tissues and promote vascular remodeling in both pregnancy and cancer^14,15^. In the adult lung, TGF-β-mediated NK cell impairment is a feature of pulmonary arterial hypertension (PAH), a disease characterized by the obstructive remodeling of the pre-capillary pulmonary arterioles.

NK cells from PAH patients exhibit excessive TGF-β signaling, cytolytic impairment and an increased production of matrix-degrading enzymes involved in vascular remodeling^16^. Moreover, mice lacking NK cells develop spontaneous pulmonary hypertension, supporting an active role for these cells in the regulation of the pulmonary circulation^17^.

Conventional NK cell functions involve the targeting of physiologically stressed, cancerous, or infected cells for cytolytic clearance. This cytotoxic activity also extends to growth-arrested senescent cells, which undergo cellular reprogramming that is marked by cell cycle arrest in response to replicative exhaustion or genotoxic stimuli^18,19^.

Although senescent cell accumulation is often recognized as a pathophysiological feature of cancer and aging^20–22^, senescence-associated cell cycle-arrest is also a central component of normal tissue patterning in multiple organs, including the developing lung^23^. Recent work has also established a role for cellular senescence in the pathogenesis of hyperoxia-induced BPD^24^, identifying a potential avenue by which NK cell senolytic activity can influence early postnatal lung development.

Here, we explore the importance of NK cell TGF-β signaling and senolytic activity to vascular and airway patterning in the neonatal lung. Through the manipulation of NK cell TGF-β signaling, we show that either excessive or insufficient clearance of senescent cells can disrupt postnatal lung development in mice. We also identify excessive NK cell TGF-β signaling and cytotoxic impairment as features of human BPD. Together, these findings present NK cell senolytic activity as a tunable regulator of lung development and a novel target for therapeutic intervention in BPD and related disorders of impaired pulmonary vascular growth.

## Materials and Methods

Detailed descriptions of experimental methods, materials, and statistical analyses are presented in the Supplemental Material.

All raw and processed sequencing data will be deposited into the Gene Expression Omibus. Code utilized to generate data will be made available on GitHub.

## Results

### Tgfbr2^NK-/-^ mice exhibit an underdeveloped pulmonary circulation and mild pulmonary hypertension

To study the impact of NK cell TGF-β signaling on the structure and function of the pulmonary circulation, we bred *Ncr1-improved Cre (iCre)^+/-^ x Ai6^+/-^ x Tgfbr2^flox/flox^* mice, hereafter referred to as *Tgfbr2^NK-/-^* mice. The presence of the *Ai6* ZsGreen reporter construct in both *Tgfbr2^NK-/-^* mice and their no-flox *Tgfbr2^NK+/+^* littermates allowed for the quantification of ZsGreen^+^ NK cell abundance in the circulation and lungs of naïve mice, which were unaltered by the Cre-mediated deletion of the TGF-βRII receptor subunit in these cells (**Fig. 1A,B**). *Ex-vivo* TGF-β treatment induced significant SMAD2 and SMAD3 phosphorylation (pSMAD2/3) in NK cells from *Tgfbr2^NK+/+^* mice, but not *Tgfbr2^NK-/-^* animals, which confirmed the insensitivity of these cells to TGF-β stimulation (**Fig. 1C**).

**Figure 1.**
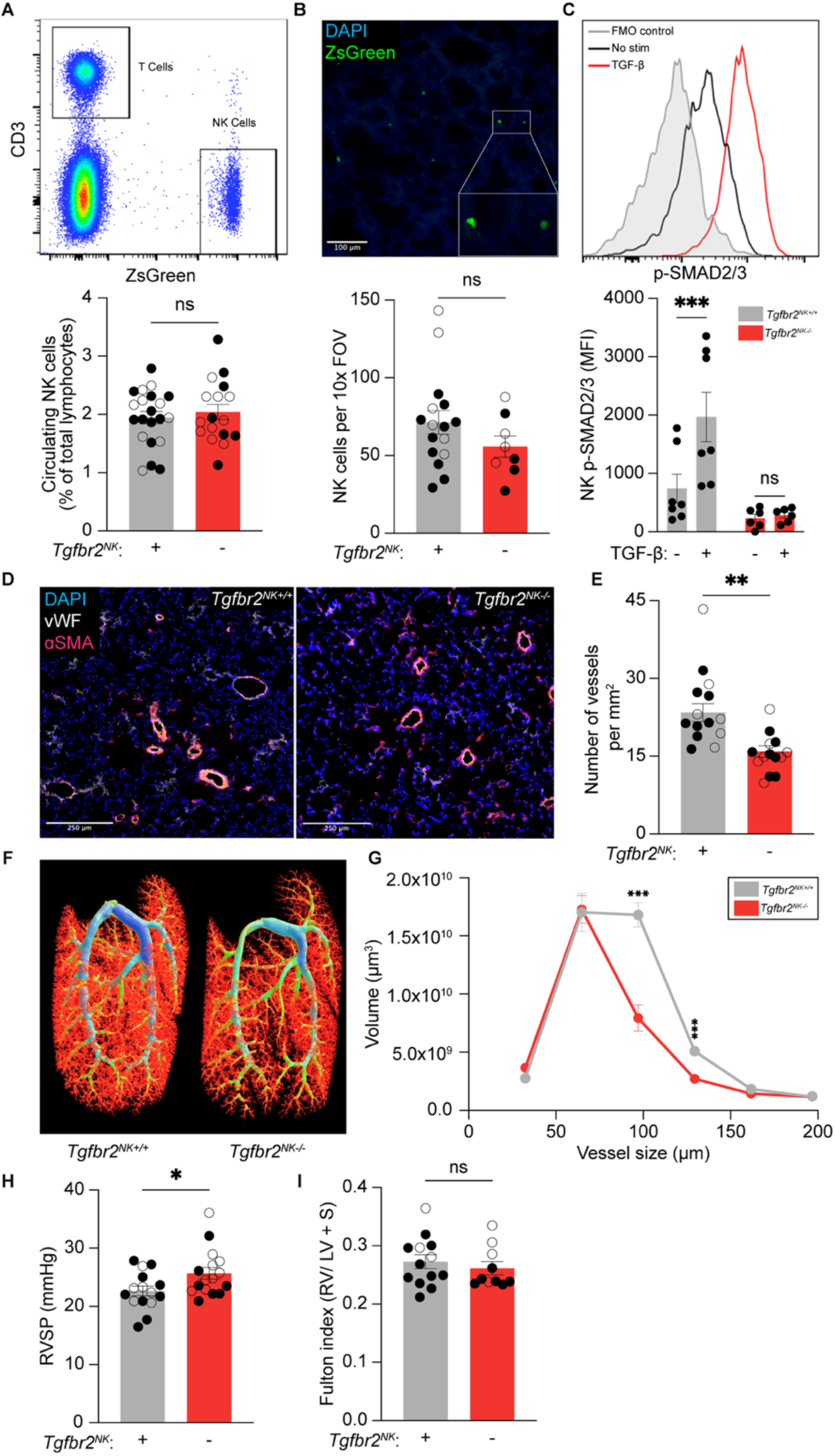
Adult *Tgfbr2^NK-/-^* mice possess a truncated pulmonary vascular tree and develop mild pulmonary hypertension. (**A**) Representative gating and quantification of circulating ZsGreen^+^/CD3^-^ NK cells in the blood of adult (8-12 week) *Tgfbr2^NK+/+^* (*n* = 21) and *Tgfbr2^NK-/-^* (*n* = 17) mice. (**B**) Quantification of ZsGreen^+^ NK cells in the lungs of adult *Tgfbr2^NK+/+^* (*n* = 16) and *Tgfbr2^NK-/-^* (*n* = 8) mice. (**C**) Representative histogram and quantification of phospho-SMAD2/3 in ZsGreen^+^ NK cells from adult *Tgfbr2^NK+/+^* (*n* = 7) and *Tgfbr2^NK-/-^* (*n* = 6) mice. (**D**) Representative images of immunofluorescent staining for von Willebrand Factor (vWF) and smooth muscle ɑ-actin (ɑSMA) in lung sections from mice of both genotypes. (**E**) Quantification of pulmonary arteriolar density in *Tgfbr2^NK+/+^* (*n* = 16) and *Tgfbr2^NK-/-^* (*n* = 16) mice. (**F**) Representative MicroCT images of the barium-perfused pulmonary vascular tree of *Tgfbr2^NK+/+^* and *Tgfbr2^NK-/-^* mice. (**G**) Assessment of pulmonary arteriolar volume in male *Tgfbr2^NK+/+^* (*n* = 3) and *Tgfbr2^NK-/-^* (*n* = 3) mice. (**H**) Assessment of right ventricular systolic pressure (RVSP) by cardiac catheterization in *Tgfbr2^NK+/+^* (*n* = 15) and *Tgfbr2^NK-/-^* (*n* = 16) mice. (**I**) Quantification of right ventricular hypertrophy (Fulton index, ratio of right ventricular (RV) weight to left ventricular (LV) weight plus septal (S) weight) for *Tgfbr2^NK+/+^* (*n* = 13) and *Tgfbr2^NK-/-^* (*n* = 10). (○) female mice, (●) male mice. ***P < 0.001, **P < 0.01, *P < 0.05, ns, not significant. Student’s t-test used in **A**-**B**, **E**, **G-I**. Two-way ANOVA, Šídák’s *post hoc* test used in **C**. Error bars are mean ± s.e.m.

Examination of the pulmonary vascular bed by the immunofluorescent quantification of pre-capillary arterioles identified a loss of distal vessels in the lungs of both male and female *Tgfbr2^NK-/-^*mice when compared to *Tgfbr2^NK+/+^* littermates (**Fig. 1D,E**). This decrease was confirmed by MicroCT imaging of barium-perfused lungs, which demonstrated a selective loss of ∼60-160 µm diameter arterioles in the lungs of *Tgfbr2^NK-/-^* mice (**Fig. 1F,G**). Importantly, this reduction was limited to arterioles of a specific size and was not observed in the very smallest pre-capillary arterioles (<60 µm) or larger vessels (>160 µm), which were equivalently abundant in animals of both genotypes. In keeping with the increased vascular resistance expected of an underdeveloped pulmonary vasculature, naïve *Tgfbr2^NK-/-^* mice exhibited mild pulmonary hypertension, as defined by an elevation in right ventricular systolic pressure (RVSP) (**Fig. 1H**). However, this modest elevation in RVSP was not accompanied by measurable right ventricular hypertrophy in *Tgfbr2^NK-/-^* mice when compared to controls (**Fig. 1I**).

### Loss of NK cell TGF-β signaling impairs postnatal alveolar development, leading to a persistent sex-dependent reduction in long-term lung function

Considering the important role for the distal pulmonary circulation in postnatal airway development, we assessed the impact of NK cell TGF-β insensitivity on alveolar patterning in the lungs of neonatal mice from postnatal day 1 (P1) to P7. Unlike humans, which undergo alveolarization from birth, mice are born with lungs at the saccular stage of development, corresponding to weeks 24-36 of human fetal patterning^42^. Assessment of distal lung morphometry by mean linear intercept (LM), a measure of the mean free distance between gas exchange surfaces, identified normal saccular airway patterning in *Tgfbr2^NK-/-^* mice from P1 to P3 (**Fig. 2A,B**). However, while *Tgfbr2^NK+/+^*controls exhibited the onset of alveolar septation and a significant decrease in LM at P5 and P7, this reduction in LM was not observed in *Tgfbr2^NK-/-^*pups, which experienced impaired alveolarization over the same period.

**Figure 2.**
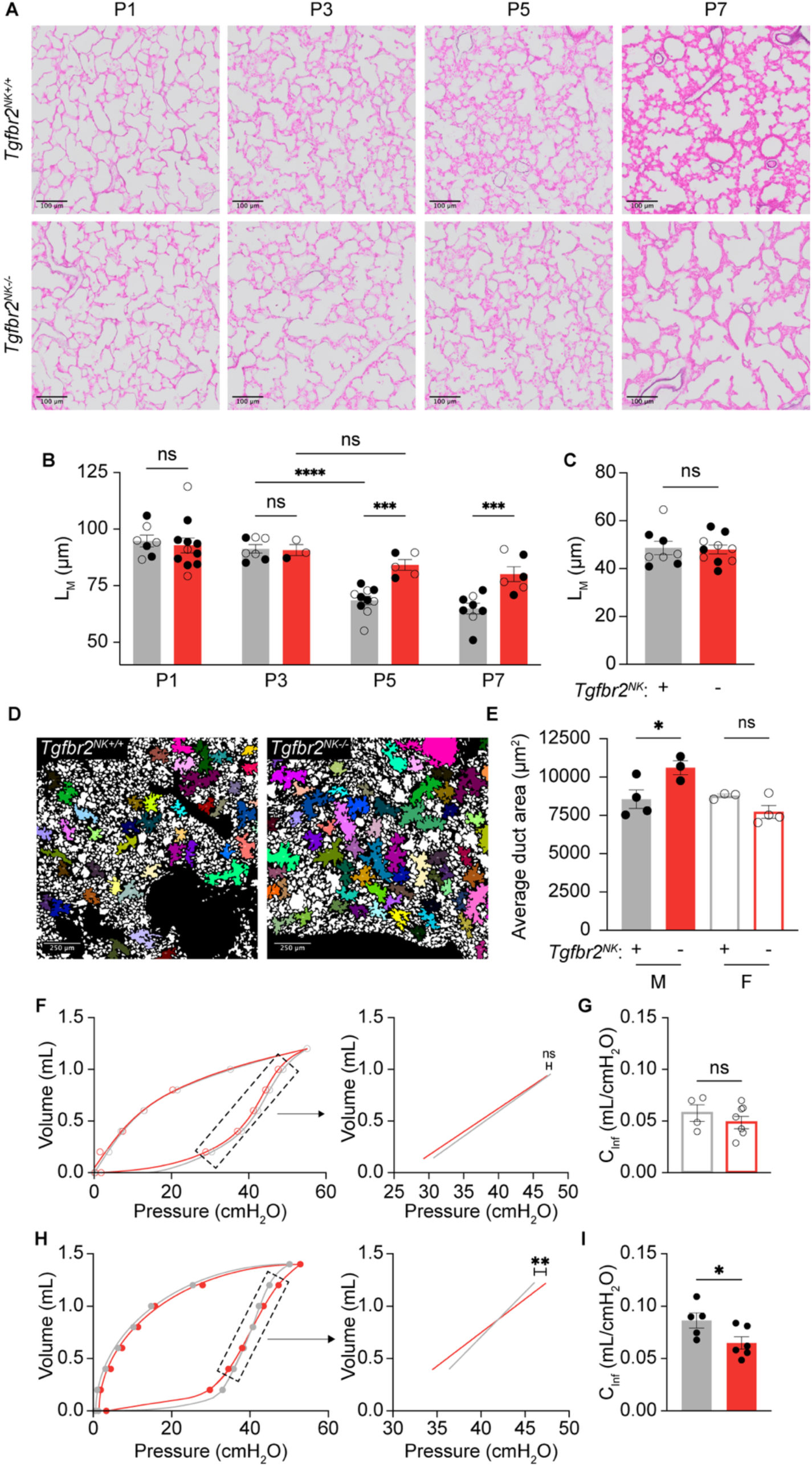
NK cell TGF-β insensitivity impairs early postnatal alveolarization and adult lung mechanics in a sex-dependent manner. (**A**) Representative images of lung sections stained using Hart’s method from mice of both genotypes at postnatal days 1 (P1), P3, P5, and P7. (**B**) Quantification of mean linear intercept (LM) at P1, P3, P5, and P7. Group sizes were: P1 *Tgfbr2^NK+/+^* (*n* = 7), P1 *Tgfbr2^NK-/-^* (*n* = 11), P3 *Tgfbr2^NK+/+^* (*n* = 7), P3 *Tgfbr2^NK-/-^* (*n* = 3), P5 *Tgfbr2^NK+/+^* (*n* = 10), P5 *Tgfbr2^NK-/-^* (*n* = 5), P7 *Tgfbr2^NK+/+^* (*n* = 8), P7 *Tgfbr2^NK-/-^* (*n* = 6). (**C**) Quantification of LM in adult *Tgfbr2^NK+/+^* (*n* = 8) and *Tgfbr2^NK-/-^* (*n* = 10) mice. (**D**) Representative binarized and masked images from adult mice of both genotypes selectively highlighting alveolar ducts. (**E**) Quantification of mean alveolar duct area in adult mice of both genotypes and sexes. Group sizes were: M *Tgfbr2^NK+/+^* (*n* = 4), M *Tgfbr2^NK-/-^* (*n* = 3), F *Tgfbr2^NK+/+^* (*n* = 3), F *Tgfbr2^NK-/-^* (*n* = 3). (**F**) Averaged pressure-volume data and visualization of inflation compliance (CInf), calculated from data bound by the dashed rectangle, from adult female *Tgfbr2^NK+/+^* (*n* = 4) and *Tgfbr2^NK-/-^* (*n* = 7) mouse lungs. (**G**) Quantification of CInf in the same group as **F**. *Tgfbr2^NK+/+^* mice are denoted by black lines, *Tgfbr2^NK-/-^* mice are denoted by red lines. (**H**) Averaged pressure-volume data and visualization of inflation compliance (CInf), calculated from data bound by the dashed rectangle from adult male *Tgfbr2^NK+/+^* (*n* = 5) and *Tgfbr2^NK-/-^* (*n* = 6) mouse lungs. (**I**) Quantification of CInf in the same group as **H**. *Tgfbr2^NK+/+^* mice are denoted by black lines, *Tgfbr2^NK-/-^* mice are denoted by red lines. (○) female mice, (●) male mice. ***P < 0.001, **P < 0.01, *P < 0.05, ns, not significant. Two-way ANOVA, Šídák’s *post hoc* test used in **B**. Student’s t-test used in **C**, **G**, **I**. One way ANOVA, Šídák’s *post hoc* test used in **E**. Simple linear regression used in **F**, **H**. Error bars are mean ± s.e.m.

Impaired alveolar septation in *Tgfbr2^NK-/-^* mice was largely resolved by adulthood, with male and female animals exhibiting normal alveolar density, lung tissue area and collagen content when compared to adult *Tgfbr2^NK+/+^* littermates (**Fig. 2C; Supplemental Fig. 1**). Assessment of intermediate respiratory airspaces did not show any differences in the overall distribution of alveolar ducts in adult *Tgfbr2^NK-/-^*mice.

However, mean duct area was significantly increased in *Tgfbr2^NK-/-^*males, but not females, when compared to sex-matched controls (**Fig. 2D,E**). Pulmonary function testing on isolated and degassed lungs confirmed a reduced inflation compliance exclusively in male *Tgfbr2^NK-/-^* mice versus *Tgrb2^NK+/+^* littermates, reflecting the higher pressure required to recruit these intermediate airspaces during inflation from a degassed state (**Fig. 2F-I**) and an overall inability of male *Tgfbr2^NK-/-^* mice to fully recover from disrupted postnatal saccular airway development.

### Impaired pulmonary vascular development in Tgfbr2^NK-/-^ mice coincides with the influx of NK cells into the neonatal lung

In keeping with the known importance of the distal pulmonary vessels in guiding postnatal airway patterning, reduced pulmonary vascularization was found to precede impaired alveolarization in *Tgfbr2^NK-/-^*mice of both sexes. Pulmonary arteriolar density was significantly decreased in the lungs of *Tgfbr2^NK-/-^* mice as early as P3, prior to the manifestation of an airway phenotype in these animals (**Fig. 3A,B**). NK cells were generally absent from the lungs of both *Tgfbr2^NK-/-^* mice and *Tgfbr2^NK+/+^*controls at birth, but numbers peaked in mice of both genotypes at P3, corresponding with the emergence of the pulmonary vascular phenotype in *Tgfbr2^NK-/-^* animals (**Fig. 3C,D**). This peak was not only significantly elevated in *Tgfbr2^NK-/-^* mice when compared to controls at P3 but remained elevated throughout the postnatal period. In contrast, lung NK cell numbers in *Tgfbr2^NK+/+^* controls returned to near-zero levels over the period from P3 to P7. Together, these findings suggest that the airway phenotype in neonatal *Tgfbr2^NK-/-^* mice arises secondary to impaired pulmonary vascular development in these animals and point to the influx of NK cells on P3 as the critical timepoint at which lung development in *Tgfbr2^NK-/-^* mice diverges from *Tgfbr2^NK+/+^*controls.

**Figure 3.**
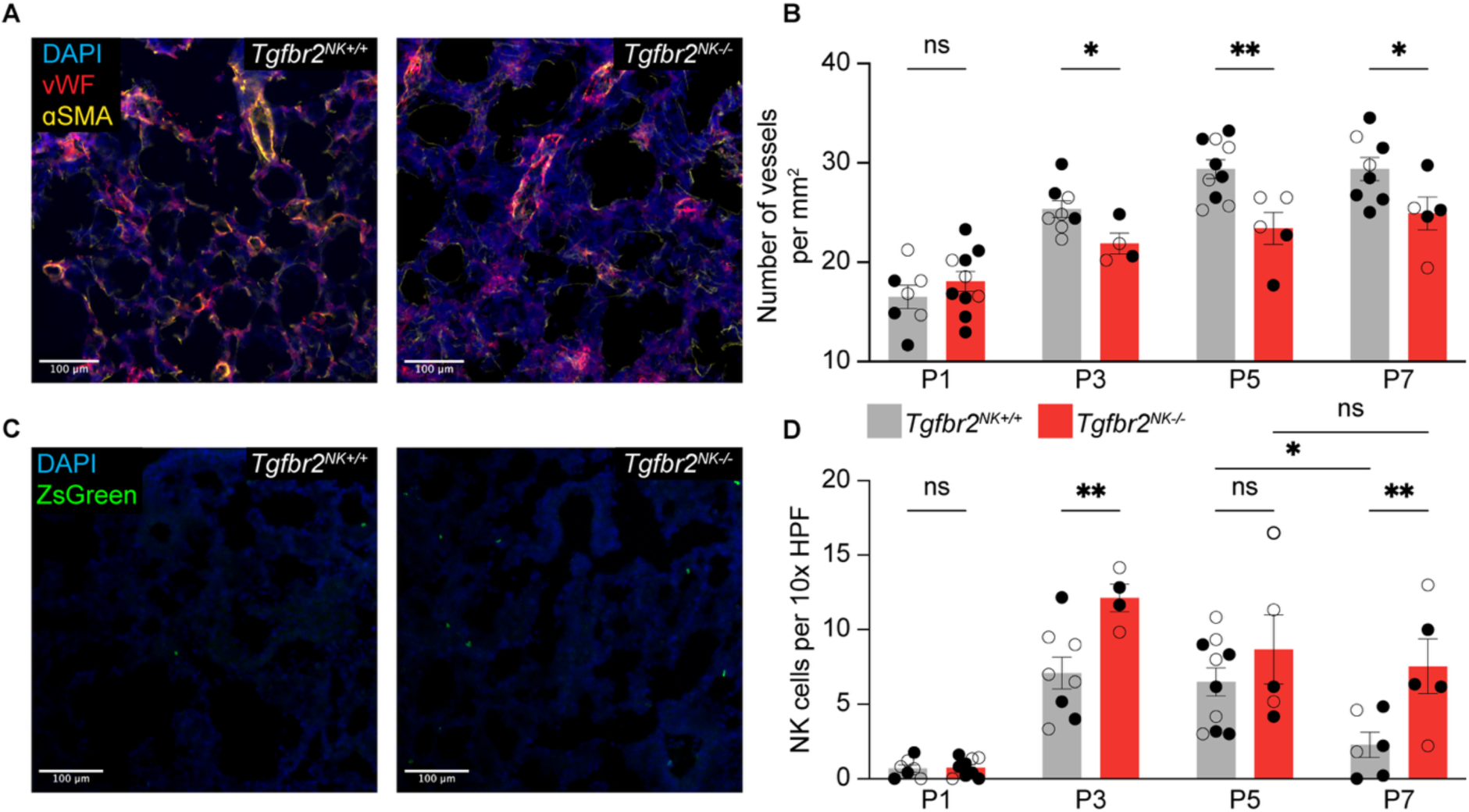
Dysregulated pulmonary vascular development coincides with the influx of NK cells into the neonatal lung at P3. (**A**) Representative images of immunofluorescent staining for vWF and ɑSMA in lung sections from mice of both genotypes at P3. (**B**) Quantification of pulmonary arteriolar density in neonatal *Tgfbr2^NK+/+^* and *Tgfbr2^NK-/-^* mice at P1, P3, P5, and P7. Group sizes were: P1 *Tgfbr2^NK+/+^* (*n* = 7), P1 *Tgfbr2^NK-/-^* (*n* = 10), P3 *Tgfbr2^NK+/+^* (*n* = 8), P3 *Tgfbr2^NK-/-^* (*n* = 4), P5 *Tgfbr2^NK+/+^* (*n* = 10), P5 *Tgfbr2^NK-/-^* (*n* = 5), P7 *Tgfbr2^NK+/+^* (*n* = 8), P7 *Tgfbr2^NK-/-^* (*n* = 5). (**C**) Representative images of ZsGreen^+^ NK cells in DAPI-stained lung sections from mice of both genotypes at P3. (**D**) Quantification of NK cell counts per 10x high-powered field (HPF) in the lungs of mice of both genotypes at P1, P3, P5, and P7. Group sizes were: P1 *Tgfbr2^NK+/+^* (*n* = 7), P1 *Tgfbr2^NK-/-^* (*n* = 9), P3 *Tgfbr2^NK+/+^* (*n* = 8), P3 *Tgfbr2^NK-/-^* (*n* = 4), P5 *Tgfbr2^NK+/+^* (*n* = 10), P5 *Tgfbr2^NK-/-^* (*n* = 5), P7 *Tgfbr2^NK+/+^* (*n* = 6), P7 *Tgfbr2^NK-/-^* (*n* = 5). (○) female mice, (●) male mice. **P < 0.01, *P < 0.05, ns, not significant. Student’s t-test used in **B** for within-timepoint comparisons. Two-way ANOVA, Šídák’s *post hoc* test used in **D** for within-and between-timepoint comparisons. Error bars are mean ± s.e.m.

### NK cell TGF-β insensitivity enhances the clearance of senescent endothelial cells in the developing lung

Having defined the pulmonary vascular and airway phenotypes of both neonatal and adult *Tgfbr2^NK-/-^* mice, single-cell RNA sequencing (scRNA-Seq) was used to assess differential gene expression in P3 lungs, with the goal of clarifying the cellular and molecular mechanisms driving impaired lung development in these animals. ScRNA-Seq on P3 lungs from *Tgfbr2^NK-/-^* mice and age-matched littermate controls allowed for the identification of broad cell groups by UMAP embedding, including lung endothelial, epithelial, mesenchymal and immune populations (**Fig. 4A; Supplemental Fig. 2A)**.

**Figure 4.**
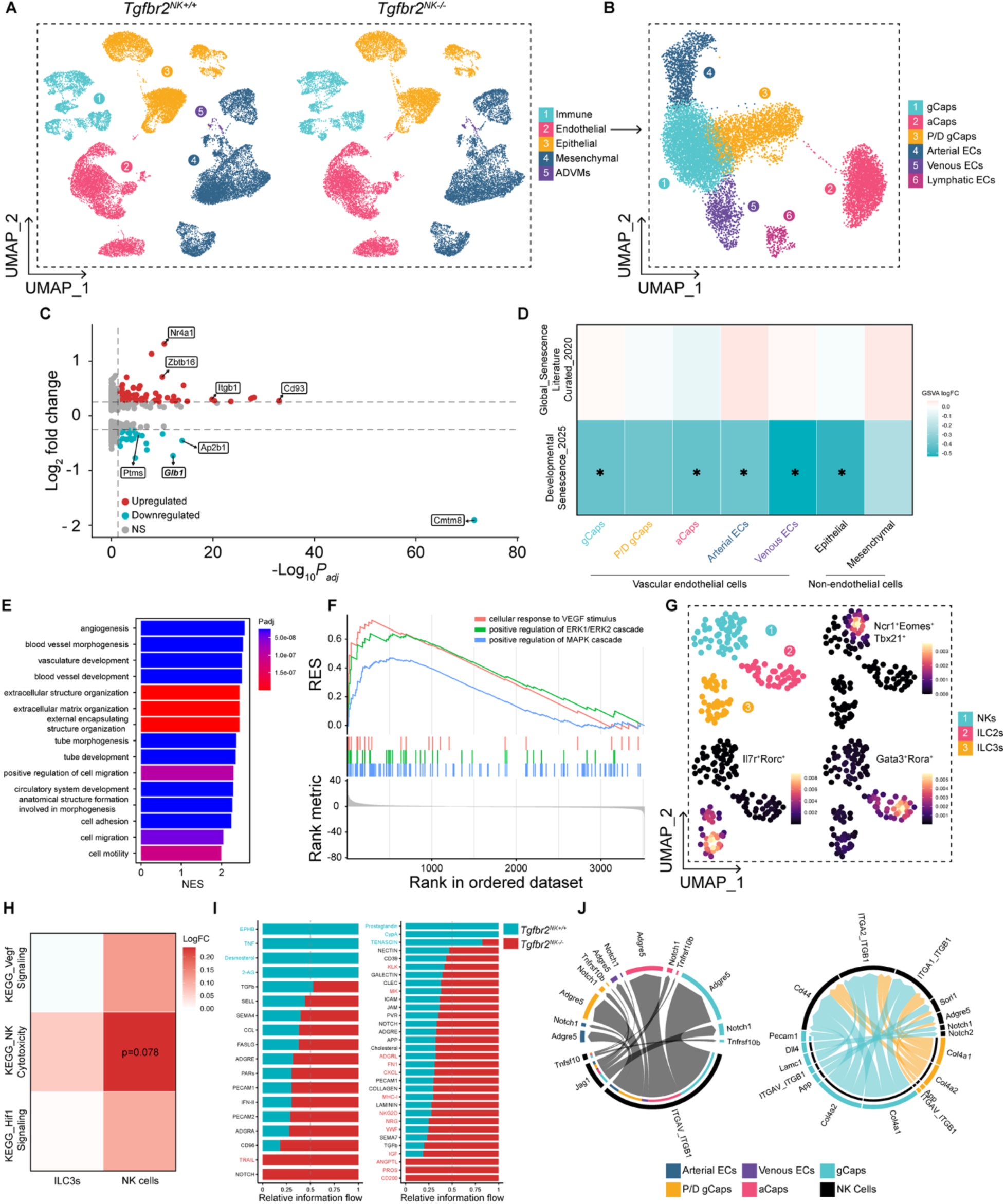
*Tgfbr2^NK-/-^* mice possess an altered NK-endothelial interactome and a lessened developmentally-associated senescence gene signature. (**A**) UMAP of all sequenced cells from neonatal *Tgfbr2^NK+/+^* (*n* = 8) and *Tgfbr2^NK-/-^* (*n* = 8) mice at P3. (**B**) UMAP of sequenced and annotated endothelial cells from both genotypes, pooled. (**C**) Volcano plot depicting genes differentially up-and down-regulated in *Tgfbr2^NK-/-^* gCap endothelial cells, relative to *Tgfbr2^NK+/+^* gCaps. (**D**) Gene set variance analysis (GSVA) matrix plot for genesets related to conventional and developmental cellular senescence. (**E**) Barplot of the top 15 enriched pathways in *Tgfbr2^NK-/-^* gCaps. (**F**) GSEA plot depicting the enrichment of three GO genesets in gCaps from *Tgfbr2^NK-/-^* mice. Red: GOBP_cellular_response_to_VEGF_stimulus, green: GOBP_positive_regulation_of_ERK1/ERK2_cascade, blue: GOBP_positive_regulation_of_MAPK_cascade. (**G**) UMAP and density plots of *Ncr1*^+^*Eomes*^+^*Tbx21*^+^ NK cells, *Gata3*^+^*Rora*^+^ ILC2s, and *Il7r*^+^*Rorc*^+^ ILC3s. (**H**) GSVA matrix plot for KEGG genesets related to NK cell cytotoxicity and proangiogenic signaling. (**I**) Stacked bar charts depicting the overall flow of information from endothelial cells to NK cells (L) and from NK cells to endothelial cells (R) for signaling pathways on the y-axis. Top signaling pathways colored blue (*Tgfbr2^NK+/+^*) or red (*Tgfbr2^NK-/-^*) are differentially enriched. (**J**) Chord diagrams depicting differentially upregulated signaling ligand-receptor pairs in *Tgfbr2^NK-/-^* mice, relative to *Tgfbr2^NK+/+^* controls, for the celltypes detailed in the legend. **P <0.01, *P < 0.05, ns, not significant. Empirical Bayes-moderated t-test used in **D,H**. Wilcoxon rank-sum test used in **I**.

Re-clustering of the endothelial cell subset enabled the delineation of lymphatic, arterial and venous endothelial populations, as well as aerocytes (aCaps) and general capillary cells (gCaps) (**Fig. 4B; Supplemental Fig. 2B**).

Differential gene expression analysis in gCaps, the population that is most likely to govern the postnatal generation of new arterioles and capillaries^43^, identified 101 differentially expressed genes in *Tgfbr2^NK-/-^* pups relative to *Tgfbr2^NK^*^+/+^ controls (**Fig. 4C**), including a downregulation of *Glb1*, the gene encoding the β-Galactosidase 1 lysosomal hydrolase (β-Gal). As a standard marker of cellular senescence^44,45^, a downregulation of β-Gal in *Tgfbr2^NK-/-^* mice raised the possibility that impaired vascular sprouting in the neonatal *Tgfbr2^NK-/-^* lung may arise as a consequence of the excessive clearance of β-Gal^+^ senescent gCaps by TGF-β-insensitive NK cells. Although gene set variation analysis (GSVA) for a broad panel of senescence-associated genes^46^ did not identify any significant changes in endothelial or non-endothelial populations from *Tgfbr2^NK-/-^* pups, a screen for a more constrained geneset of markers previously detected in developmentally-associated senescent cells^24,47^ identified the downregulation of this geneset in several endothelial cell subsets, including gCaps and arterial endothelial cells, as well the pulmonary epithelium of *Tgfbr2^NK-/-^* mice (**Fig. 4D**). Furthermore, gene set enrichment analysis (GSEA) identified the upregulation of several biological pathways associated with angiogenesis, blood vessel formation and tube formation in *Tgfbr2^NK-/-^* gCaps (**Fig. 4E**). This was accompanied by the positive enrichment of genesets relating to ERK-MAPK signaling and VEGF stimulation (**Fig. 4F**). Together, these findings suggest the upregulation of a proangiogenic, potentially compensatory, transcriptional program in gCaps driven by VEGF signaling^48^ in response to the excessive pruning and inadequate pulmonary vascular development in *Tgfbr2^NK-/-^* mice.

To explore the impact of TGF-β insensitivity on NK cells in the developing lung, re-clustering was performed on immune cells to identify NKs for further analysis (**Fig. 4G; Supplemental Fig. 2C**). GSVA revealed the upregulation of the KEGG NK-mediated cytotoxicity pathway in NK cells from *Tgfbr2^NK-/-^* mice, (**Fig. 4H**), with no significant changes in hypoxia-associated pathways linked to angiogenesis and vascular remodeling. A similar examination of ILC3s, a lymphocyte subset which can express Nkp46 and adopt a cytotoxic phenotype, identified no differences in cytotoxicity or hypoxia-associated pathways in these cells. Examination of NK-vascular crosstalk identified the upregulation of outgoing TRAIL signaling from NK cells to endothelial cells in *Tgfbr2^NK-/-^* lungs, a signaling pathway linked to NK cell cytotoxicity.^49^ (**Fig. I**).

Interestingly, outgoing ephrin-B (EPHB) signaling from NK cells to endothelial cells was absent in *Tgfbr2^NK-/-^*mice. EPHB signaling, specifically through the ephrin-B2 receptor, has been previously reported as a critical contributor to uterine NK cell-driven angiogenesis and endothelial tube formation^50^, suggesting reductions in pulmonary vascular density may result in part from the reduced angiogenic capacity of lung NK cells in *Tgfbr2^NK-/-^* mice. The heightened engagement of NK cells without the type-II TGF-β receptor was further supported by the identification of increased incoming NKG2D signaling from endothelial cells (**Fig. 4I**), outgoing Tnfsf10 signaling to several endothelial cell populations, and incoming Dll4 signaling^51^ from gCaps (**Fig. 4J**).

Validation of the scRNA-Seq findings by assessment of senescence-associated (SA)-β-Gal activity revealed a significant reduction of SA-β-Gal^+^ cells in the lungs of P3 *Tgfbr2^NK-/-^*mice relative to age-matched controls (**Fig. 5A,B**), validating the findings from the scRNA-Seq analysis. Immunofluorescent staining for endomucin (Emcn), a marker of capillary-associated endothelial cells^52^, confirmed the presence of β-Gal^+^ capillary cells in the lung at P3 (**Fig. 5C**). The enhanced capacity of NK cells from *Tgfbr2^NK-/-^* mice to lyse senescent fetal mouse lung endothelial cells was also validated by *ex-vivo* co-culture, with NK cells from these animals exhibiting both an increased baseline senolytic activity and insensitivity to TGF-β mediated suppression when compared to NK cells from *Tgfbr2^NK^*^+/+^ controls (**Fig. 5D,E**).

**Figure 5.**
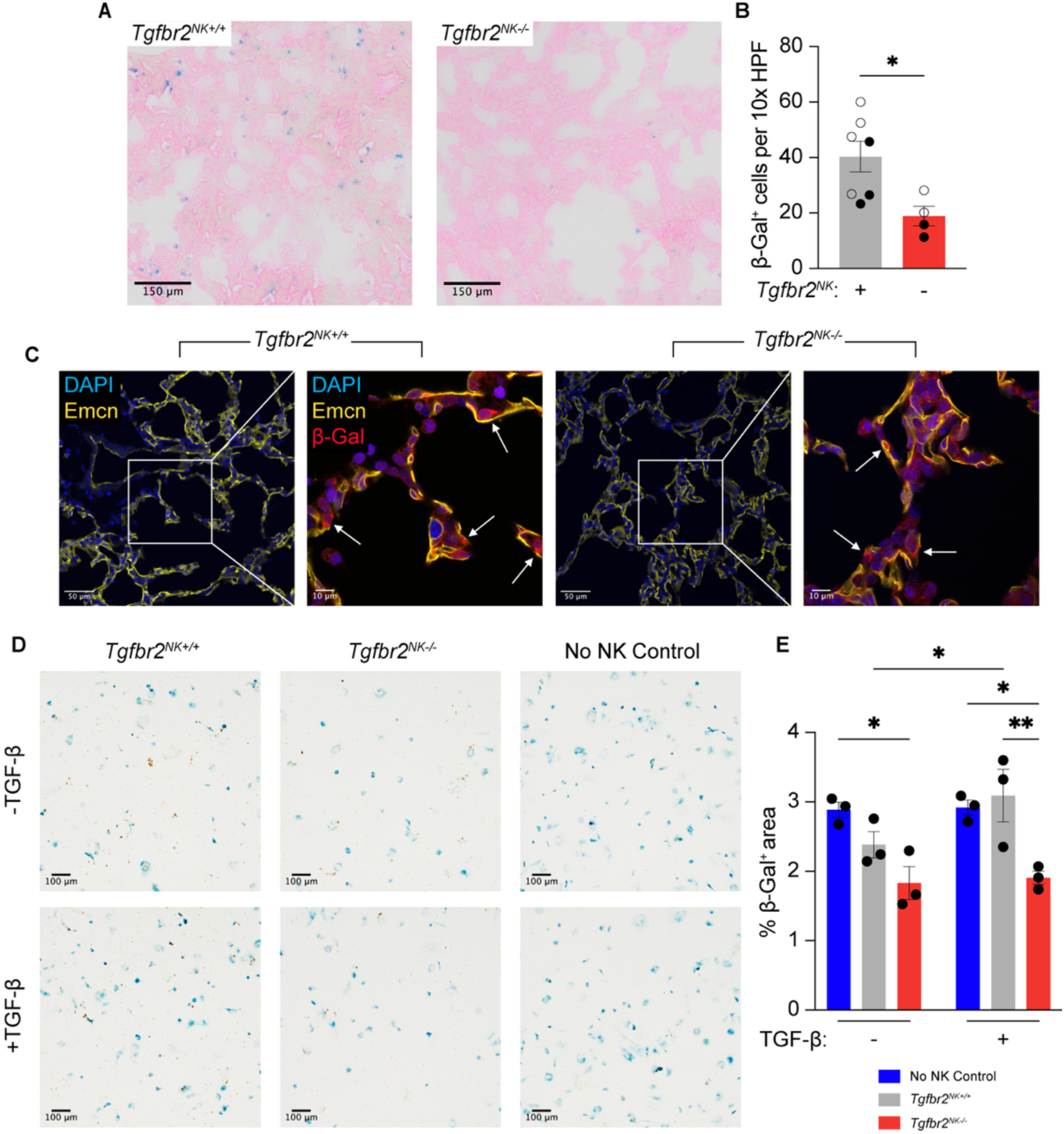
NK cell TGF-β insensitivity enhances the clearance of β-galactosidase^+^ senescent endothelial cells in the developing lung. (**A**) Representative images of immunohistochemical staining for senescence-associated β-galactosidase (β-gal) in lung sections from mice of both genotypes at P3. (**B**) Quantification of the number of β-Gal^+^ cells per 10x HPF in neonatal *Tgfbr2^NK+/+^* (*n* = 7) and *Tgfbr2^NK-/-^* (*n* = 4) mice at P3. (**C**) Representative images of immunofluorescent staining for Emcn and β-Gal in lung sections *from Tgfbr2^NK+/+^* and *Tgfbr2^NK-/-^* mice at P3. White arrows depict β-Gal-expressing senescent cells. (**D**) Representative images of immunohistochemical staining for SA-β-Gal in MFLM-91U cells following 8-hour co-culture. (**E**) Quantification of percent β-Gal^+^ area per condition. Group sizes were (*n* = 3) for all conditions. (○) female mice, (●) male mice. **P <0.01, *P < 0.05, ns, not significant. Student’s t-test used in **B**. Two-way ANOVA, Šídák’s *post hoc* test used in **E**. Error bars are mean ± s.e.m.

### NK cell TGF-β signaling is elevated in human BPD

The identification of enhanced senescent cell clearance in the lungs of *Tgfbr2^NK-/-^* mice directly contrasts with models of hyperoxia-induced BPD, for which impaired lung patterning has been linked to elevated TGF-β signaling and an accumulation of senescent cells^6,24^. Moreover, assessment of the pulmonary circulation in adult *Nfil3^-/-^* mice, a strain that is characterized by NK cell deficiency, not hyperactivity, identified a reduction in pulmonary arteriolar density (**Supplemental Fig. 3**) that mirrored the decrease seen in *Tgfbr2^NK-/-^* animals. In order to address these seemingly contradictory findings and determine the clinical relevance of NK cell TGF-β signaling to human BPD, scRNA-Seq data was analyzed from human infants with BPD at term-corrected gestational age and age-matched controls^53^. Clustering and embedding of lung immune cell subsets allowed for the identification of NK cells, which were significantly reduced in proportion in the BPD samples as a proportion of total immune cells (**Fig. 6A,B**). GSEA, performed using a ranked gene list derived exclusively from tested genes in NK cells, identified several genesets related to effector functionalities that were negatively enriched in the samples from BPD infants (**Fig. 6C**). This reduction was accompanied by the positive enrichment of genes involved in the cellular response to TGF-β, supporting a link between heightened TGF-β signaling and impaired NK cell function in disease.

**Figure 6.**
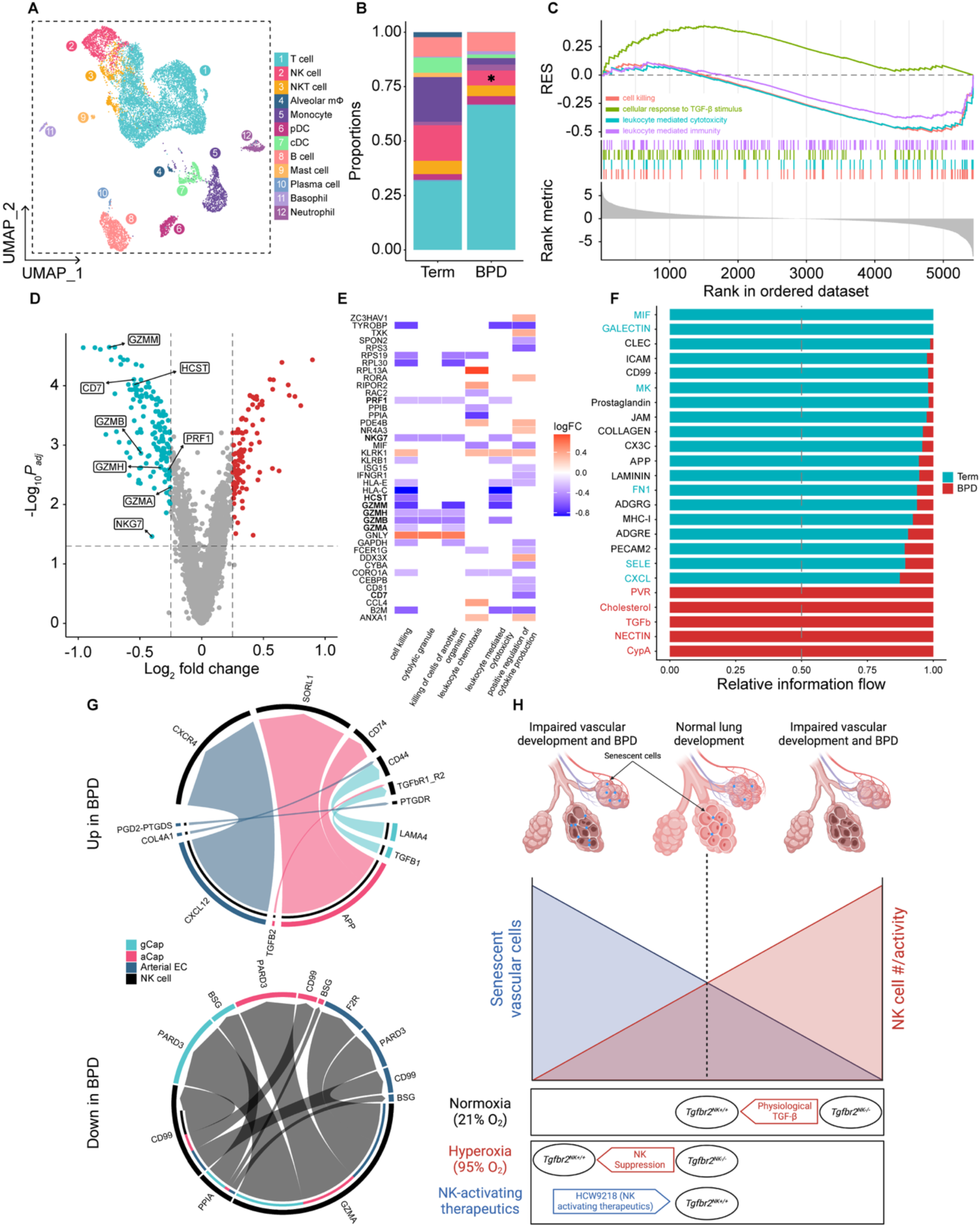
NK cell dysfunction is a feature of BPD in human infants. (**A**) UMAP plot of sequenced and labelled immune cells, pooled from human infants with BPD (*n* = 2) and term controls (*n* = 2). (**B**) Stacked bar plot of cellular proportions binned by condition. Color corresponds to the respective celltype in **A**. (**C**) GSEA plot depicting the enrichment of four GO genesets in NK cells from infants with BPD. Red: GOBP_cell_killing, green: GOBP_cellular_response_to_TGF-β _stimulus, blue: GOBP_leukocyte_mediated_cytotoxicity, purple: GOBP_leukocyte_mediated_immunity. (**D**) Volcano plot of differentially expressed genes in NK cells from infants with BPD relative to term controls. (**E**) Heatplot of up-and down-regulated genes from significantly overrepresented genesets in NK cells from infants with BPD. Bolded genes are labelled in (**D**). (**F**) Stacked bar chart depicting the overall flow of information from endothelial cells to NK cells for signaling pathways on the y-axis. Top signaling pathways colored blue (term controls) or red (BPD) are differentially enriched. (**G**) Chord diagrams depicting upregulated and downregulated ligand-receptor pairings in NK cells and endothelial cells from infants with BPD, relative to term controls. (**H**) Proposed model depicting the actions of NK cell activity and cellular senescence on early postnatal lung patterning. *P < 0.05. Wilcoxon rank-sum test used in **F**.

Differential expression analysis identified the downregulation of several key genes associated with NK cell cytotoxic function in BPD, including 4 of the 5 human granzymes expressed by NK cells (**Fig. 6D**). Of note, many of the downregulated genes in BPD NK cells belonged to cytotoxic-and immune response-related gene ontology (GO) genesets that were significantly overrepresented in disease (**Fig. 6E**), further supporting NK cell functional impairment as a feature of human BPD. CellChat analysis between NK cells and the pulmonary endothelium identified an upregulation of outgoing TGF-β signaling from aCap and gCap endothelial populations to NK cells in BPD, alongside a downregulation of outgoing GZMA ligand-receptor pairings from NK cells to the endothelium (**Fig. 6F,G**). When coupled with our findings from the *Tgfbr2^NK-/-^*mice, these results suggest that NK cell TGF-β signaling may serve as a tunable regulator of postnatal lung patterning. Under this model, any change in NK cell activity in the lung, either linked to TGF-β insensitivity and enhanced senescent cell clearance (*Tgfbr2^NK-/-^* mice), arising secondary to TGF-β mediated NK cell impairment (human BPD, hyperoxia models) or as a consequence of absolute NK cell loss (*Nfil3^-/-^* mice), would lead to the disruption of pulmonary vascular patterning and airway development (**Fig. 6H**).

### NK cell TGF-β-insensitivity confers partial protection from hyperoxia-induced BPD

The identification of TGF-β-mediated NK cell suppression in human BPD raised the possibility that targeting the NK cell-TGF-β signaling axis may provide a means of ameliorating disease by restoring cytotoxic function and enhancing the clearance of pathological senescent cells. To explore this hypothesis, we exposed newborn *Tgfbr2^NK+/+^* and *Tgfbr2^NK-/-^* mice at P0 to 95% O2 for 72 hours as an acute model of hyperoxia-induced BPD. O2 exposure from P0 to P3 resulted in impaired alveolar development in *Tgfbr2^NK+/+^* mice, relative to normoxic wild-type controls (**Fig. 7A,B**), in addition to reductions in pulmonary arteriolar density (**Fig. 7C,D**). Interestingly, *Tgfbr2^NK-/-^* mice were partially protected from these impairments in alveolar and pulmonary vascular development, supporting the hypothesis that NK cell TGF-β signaling may be a relevant target in hyperoxia-induced BPD.

**Figure 7.**
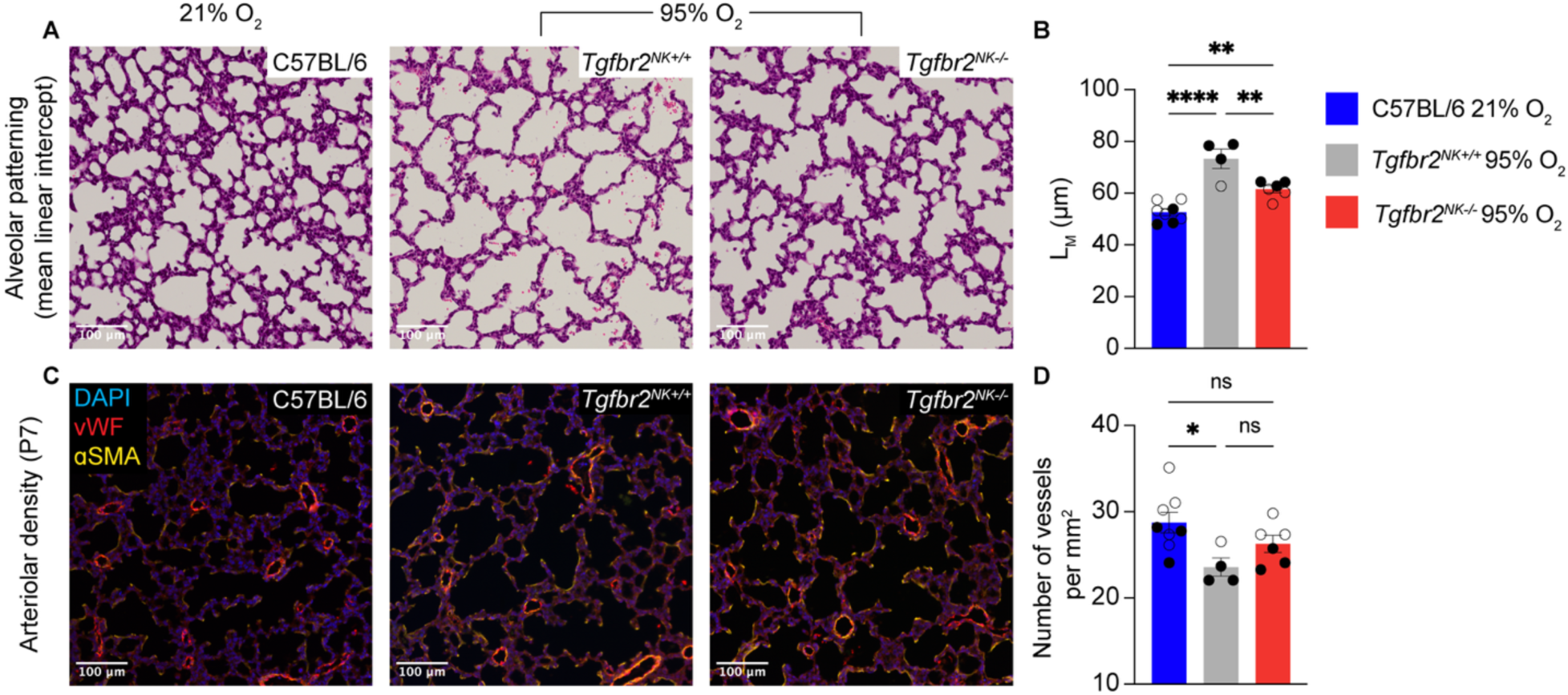
Neonatal *Tgfbr2^NK-/-^* mice are partially protected from hyperoxia-induced BPD. (**A**) Representative images of immunofluorescent staining for vWF and ɑSMA in lung sections from wild-type and *Tgfbr2^NK^* neonatal mice at P7 following exposure to normoxia (21% O2) or hyperoxia (95% O2) for 3 days. Group sizes were: 21% C57BL/6 WT (*n* = 8), 95% *Tgfbr2^NK+/+^* (*n* = 4), 95% *Tgfbr2^NK-/-^* (*n* = 6). (**B**) Quantification of pulmonary arteriolar density in the same groups as in **A**. (**C**) Representative images of immunohistochemical hematoxylin and eosin staining in lung sections from mice in **A**. (**D**) Quantification of LM in the same groups as in **A**. (○) female mice, (●) male mice. ****P < 0.0001, **P < 0.01, *P < 0.05, ns, not significant. One-way ANOVA, Tukey’s *post hoc* test used in **B**, **D**. Error bars are mean ± s.e.m.

### Prevention of hyperoxia-induced BPD using an NK cell-targeted senolytic

To test the viability of targeting excessive TGF-β signaling and NK cell impairment as a therapeutic avenue for BPD, we explored the potential of HCW9218, an NK cell-activating senolytic consisting of a bifunctional TGF-β ligand trap fused to an IL-15/IL-15 receptor-ɑ superagonist^54^ (**Fig. 8A**), to rescue lung development in an acute hyperoxia-induced model of BPD (**Fig. 8B**). Flow cytometry confirmed that HCW9218 induced an expansion of NK cells in both the circulation (**Fig. 8C**) and lungs (**Fig. 8D**) of treated animals by P3 relative to vehicle-only 21% and 95% O2 controls. Additionally, a greater proportion of these expanded cells were positive for granzyme B (Gzmb; **Fig. 8E,F,G**). Interestingly, a reduced proportion of circulating NK cells from vehicle-only 95% O2 controls were Gzmb^+^ relative to 21% O2 controls, suggesting that hyperoxia-induced NK cell dysfunction may extend beyond the pulmonary circulation.

**Figure 8.**
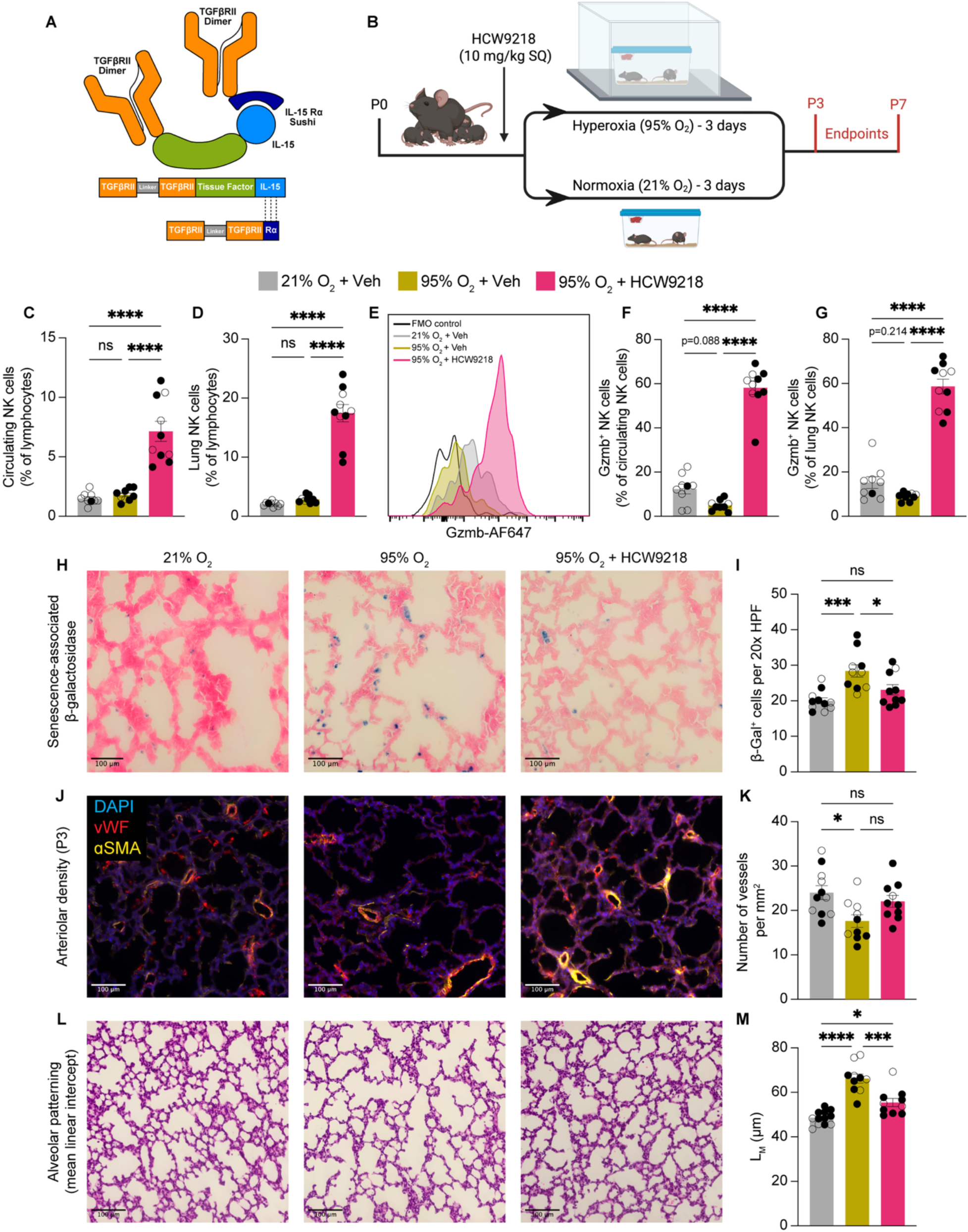
Activation of NK cells by HCW9218 attenuates hyperoxia-induced BPD. (**A**) Graphical representation of HCW9218. (**B**) Schematic of the utilized experimental timeline for testing. (**C**) Quantification of CD45^+^NK1.1^+^CD3^-^ NK cells in the circulationof neonatal mice at P7 following exposure to either normoxia (21% O2) or hyperoxia (95% O2) for 3 days, expressed as a percent of total lymphocytes. Group sizes were: P7 21% vehicle (*n* = 9), P7 95% vehicle (*n* = 9), P7 95% HCW9218 (*n* = 10) (**D**) Quantification of CD45^+^NK1.1^+^CD3^-^ NK cells in the lungs of the neonatal mice in **C**, expressed as a percent of total lymphocytes. (**E**) Representative histogram of Gzmb expression in CD45^+^NK1.1^+^CD3^-^ NK cells (**F**) Quantification of circulating Gzmb^+^ NK cells in neonatal mice from **C**, expressed as a percent of total NK cells. (**G**) Quantification of Gzmb^+^ NK cells in the lungs of the neonatal mice in **C**, expressed as a percent of total NK cells. (**H**) Representative images of immunohistochemical staining for β-Gal^+^ cells in lung sections from mice at P3 given vehicle or HCW9218 (10 mg/kg, S.Q.) and exposed to either normoxia (21% O2) or hyperoxia (95% O2) for 3 days. Group sizes were: P3 21% vehicle (*n* = 11), P3 95% vehicle (*n* = 10), P3 95% HCW9218 (*n* = 10). (**I**) Quantification of β-Gal^+^ cells per 20x HPF in the same groups as in **H**. (**J**) Representative images of immunofluorescent staining for vWF and ɑSMA in lung sections from neonatal mice in **H**. (**K**) Quantification of pulmonary arteriolar density in the same groups as in **H**. (**L**) Representative images of immunohistochemical hematoxylin and eosin staining in lung sections from wild-type mice dosed with vehicle or HCW9218 at P7. Group sizes were: P7 21% vehicle (*n* = 12), P7 95% vehicle (*n* = 10), P7 95% HCW9218 (*n* = 10). (**M**) Quantification of LM in the same groups as in **L**. (○) female mice, (●) male mice. ****P < 0.0001, ***P < 0.001, *P < 0.05, ns, not significant. One-way ANOVA, Tukey’s *post hoc* test used in **C**, **D**, **F, G, I, K, M**. Error bars are mean ± s.e.m.

Exposure of neonatal mice to 95% O2 for 72 hours from P0 to P3 induced an accumulation of β-Gal^+^ senescent cells in the lungs (**Fig. 8H,I**), which was accompanied by decreased pulmonary vascular density at P3 (**Fig. 8J,K**) and impaired alveolarization at P7 (**Fig. 8L,M**). Treatment of pups with 10 mg/kg of HCW9218 immediately prior to hyperoxia exposure on P0 not only prevented the accumulation of β-Gal^+^ cells in the lung but also protected mice from the impaired vascular and airway development seen in vehicle-treated controls, indicating that the promotion of immune-mediated senolytic activity can be beneficial in this model.

## Discussion

NK cell subsets have been shown to significantly influence vascular structure and patterning in cases of physiological remodeling, like pregnancy^55^, and in disease states like cancer, where infiltrating NK cells contribute to tumor angiogenesis through the release of proangiogenic factors^15^. However, the extension of this functionality to the development and maintenance of the pulmonary circulation had not yet been explored. By experimentally targeting TGF-β signaling in NK cells, we have identified a new role for this immune cell population as a critical regulator of late-stage saccular lung development and a potential target for therapeutic intervention in BPD. While the identification of reduced pulmonary vascular density in the lungs of *Tgfbr2^NK-/-^* mice shows that some degree of physiologic TGF-β signaling is required to temper NK cell activity and guide normal patterning, our work in the hyperoxia-induced BPD model demonstrates that TGF-β sequestration and the promotion of IL-15 signaling can also serve to restore alveolarization and vascular development in cases where TGF-β signaling is excessive. This capacity of TGF-β to serve as a tunable regulator of NK cell activity, senescent cell clearance and postnatal lung patterning makes this pathway an appealing target for therapeutic intervention in diseases of impaired lung development.

Importantly, similar pulmonary vascular defects were identified in the lungs of both NK cell-deficient *Nfil3^-/-^* mice and *Tgfbr2^NK-/-^*animals, validating the critical role for NK cells in distal lung patterning in two independent genetic models. The identification of elevated TGF-β signaling and reduced cytotoxic gene expression in NK cells from infants with BPD by single cell RNA sequencing not only supports these findings from the mouse models, but also provides the first reported evidence of NK cell dysfunction as a feature of the human disease. Additionally, by demonstrating that the genetic insensitivity of NK cells to TGF-β signaling in neonatal *Tgfbr2^NK-/-^*mice confers partial protection against disease, we have provided independent validation for the finding that NK cell dysfunction as a result of elevated TGF-β signaling is a contributor to hyperoxia-induced BPD.

Although our previous work had reported the development of spontaneous pulmonary hypertension in aged *Nfil3*^-/-^ mice^17^, the link between the adult cardiopulmonary phenotype and impaired postnatal lung development was not assessed in that study. It is noteworthy that antenatal lung development appears to be unaltered in *Tgfbr2^NK-/-^*mice, with differences only arising alongside the transient influx of NK cells into the lung during the early postnatal period. This timeline differs from other conditional mouse models of tissue-specific *Tgfbr2* deletion, which involve disrupted lung branching prior to birth in mesenchymal, but not epithelial *Tgfbr2^-/-^* mice^56,57^. These varied phenotypes arising from the cell type-specific deletion of *Tgfbr2* highlight the complex and tissue-dependent contributions of TGF-β signaling to specific stages of pre-and postnatal lung development.

The effects of NK cell TGF-β insensitivity on postnatal vascular and airway development were seen in both male and female *Tgfbr2^NK-/-^* neonates. However, with the exception of the reduction in pulmonary arteriolar density, many of the differences in lung structure and function persisted only in *Tgfbr2^NK-/-^* males. This persistence reflects a clinical reality, whereby male preterm infants are at a higher risk for adverse outcomes, including BPD^58,59^, but differs from the adult human population, where female survivors of BPD are more susceptible to developing impaired pulmonary function later in life^60^.

Additional studies are required to determine the cellular and molecular processes governing catch-up airway development in female, but not male *Tgfbr2^NK-/-^* mice.

Our examination of HCW9218 in the hyperoxic model of BPD offers a proof-of-concept for this Phase I approved senolytic (Trial Identifiers: NCT05304936, NTC05322408) in a disease model that mimics the clinical reality of oxygen supplementation in extreme prematurity. The current study also offers new insights into the emerging role for senescent cells as active contributors to tissue patterning across a range of organs.

This recognition that senescent cells not only contribute to disease pathology, but are also essential to tissue development and homeostasis, highlights a major consideration in the translation of senolytics as therapeutics for a broad array of diseases. In this context, immune-mediated senolytic clearance with compounds like HCW9218 may offer an advantage over small molecule senolytics targeting anti-apoptotic mediators^61^ by promoting the selective removal of pathological senescent cells, while leaving those required for normal tissue patterning and function undisturbed. Future work exploring this therapeutic strategy may involve the validation of HCW9218 in BPD models driven by longer-term hyperoxia exposure^62^, as well as studies examining the effect of treatment on long-term lung function and cardiopulmonary haemodynamics into adulthood.

## Acknowledgements

We would like to thank Emilie Ward and Jenny Thiele for their technical assistance with the lung functional testing, and to Professor Emeritus Dr. John Fisher for having introduced this technique to NJD.

## Supplemental Material

Extended Methods

Tables S1-S4

Figures S1-S3

References 25-41

## References

1. Schittny JC. Development of the lung. Cell and Tissue Research. 2017;367:427–444. doi: 10.1007/s00441-016-2545-0

2. Le Cras TD, Rabinovitch M. Pulmonary Vascular Development. In: Jobe AH, Whitsett JA, Abman SH, eds. Fetal and Neonatal Lung Development: Clinical Correlates and Technologies for the Future. Cambridge: Cambridge University Press; 2016:34-57.

3. Jobe AJ. The New BPD: An Arrest of Lung Development. Pediatric Research. 1999;46:641–641. doi: 10.1203/00006450-199912000-00007

4. D’Angio CT, Maniscalco WM. The role of vascular growth factors in hyperoxia-induced injury to the developing lung. Front Biosci. 2002;7:d1609–1623. doi: 10.2741/a865

5. Thébaud B, Goss KN, Laughon M, Whitsett JA, Abman SH, Steinhorn RH, Aschner JL, Davis PG, McGrath-Morrow SA, Soll RF, et al. Bronchopulmonary dysplasia. Nature Reviews Disease Primers. 2019;5:78. doi: 10.1038/s41572-019-0127-7

6. Calthorpe RJ, Poulter C, Smyth AR, Sharkey D, Bhatt J, Jenkins G, Tatler AL. Complex roles of TGF-β signaling pathways in lung development and bronchopulmonary dysplasia. American Journal of Physiology-Lung Cellular and Molecular Physiology. 2023;324:L285–L296. doi: 10.1152/ajplung.00106.2021

7. Hilgendorff A, Parai K, Ertsey R, Jain N, Navarro EF, Peterson JL, Tamosiuniene R, Nicolls MR, Starcher BC, Rabinovitch M, et al. Inhibiting Lung Elastase Activity Enables Lung Growth in Mechanically Ventilated Newborn Mice. American Journal of Respiratory and Critical Care Medicine. 2011;184:537–546. doi: 10.1164/rccm.201012-2010OC

8. Mokres LM, Parai K, Hilgendorff A, Ertsey R, Alvira CM, Rabinovitch M, Bland RD. Prolonged mechanical ventilation with air induces apoptosis and causes failure of alveolar septation and angiogenesis in lungs of newborn mice. American Journal of Physiology-Lung Cellular and Molecular Physiology. 2010;298:L23–L35. doi: 10.1152/ajplung.00251.2009

9. Massagué J, Sheppard D. TGF-β signaling in health and disease. Cell. 2023;186:4007–4037. doi: 10.1016/j.cell.2023.07.036

10. Saito A, Horie M, Nagase T. TGF-β Signaling in Lung Health and Disease. Int J Mol Sci. 2018;19. doi: 10.3390/ijms19082460

11. Chen H, Zhuang F, Liu YH, Xu B, Del Moral P, Deng W, Chai Y, Kolb M, Gauldie J, Warburton D, et al. TGF-beta receptor II in epithelia versus mesenchyme plays distinct roles in the developing lung. Eur Respir J. 2008;32:285–295. doi: 10.1183/09031936.00165407

12. Miao Q, Chen H, Luo Y, Chiu J, Chu L, Thornton ME, Grubbs BH, Kolb M, Lou J, Shi W. Abrogation of mesenchyme-specific TGF-β signaling results in lung malformation with prenatal pulmonary cysts in mice. Am J Physiol Lung Cell Mol Physiol. 2021;320:L1158–l1168. doi: 10.1152/ajplung.00299.2020

13. Viel S, Marcais A, Guimaraes FS, Loftus R, Rabilloud J, Grau M, Degouve S, Djebali S, Sanlaville A, Charrier E, et al. TGF-beta inhibits the activation and functions of NK cells by repressing the mTOR pathway. Sci Signal. 2016;9:ra19. doi: 10.1126/scisignal.aad1884

14. Ratsep MT, Felker AM, Kay VR, Tolusso L, Hofmann AP, Croy BA. Uterine natural killer cells: supervisors of vasculature construction in early decidua basalis. Reproduction. 2015;149:R91–102. doi: 10.1530/REP-14-0271

15. Gotthardt D, Putz EM, Grundschober E, Prchal-Murphy M, Straka E, Kudweis P, Heller G, Bago-Horvath Z, Witalisz-Siepracka A, Cumaraswamy AA, et al. STAT5 Is a Key Regulator in NK Cells and Acts as a Molecular Switch from Tumor Surveillance to Tumor Promotion. Cancer Discovery. 2016;6:414–429. doi: 10.1158/2159-8290.Cd-15-0732

16. Ormiston ML, Chang C, Long LL, Soon E, Jones D, Machado R, Treacy C, Toshner MR, Campbell K, Riding A, et al. Impaired natural killer cell phenotype and function in idiopathic and heritable pulmonary arterial hypertension. Circulation. 2012;126:1099–1109. doi: 10.1161/CIRCULATIONAHA.112.110619

17. Ratsep MT, Moore SD, Jafri S, Mitchell M, Brady HJM, Mandelboim O, Southwood M, Morrell NW, Colucci F, Ormiston ML. Spontaneous pulmonary hypertension in genetic mouse models of natural killer cell deficiency. Am J Physiol Lung Cell Mol Physiol. 2018;315:L977–L990. doi: 10.1152/ajplung.00477.2017

18. Xue W, Zender L, Miething C, Dickins RA, Hernando E, Krizhanovsky V, Cordon-Cardo C, Lowe SW. Senescence and tumour clearance is triggered by p53 restoration in murine liver carcinomas. Nature. 2007;445:656–660. doi: 10.1038/nature05529

19. Brighton PJ, Maruyama Y, Fishwick K, Vrljicak P, Tewary S, Fujihara R, Muter J, Lucas ES, Yamada T, Woods L, et al. Clearance of senescent decidual cells by uterine natural killer cells in cycling human endometrium. eLife. 2017;6:e31274. doi: 10.7554/eLife.31274

20. Baker DJ, Perez-Terzic C, Jin F, Pitel KS, Niederländer NJ, Jeganathan K, Yamada S, Reyes S, Rowe L, Hiddinga HJ, et al. Opposing roles for p16Ink4a and p19Arf in senescence and ageing caused by BubR1 insufficiency. Nat Cell Biol. 2008;10:825–836. doi: 10.1038/ncb1744

21. Coppé JP, Patil CK, Rodier F, Sun Y, Muñoz DP, Goldstein J, Nelson PS, Desprez PY, Campisi J. Senescence-associated secretory phenotypes reveal cell-nonautonomous functions of oncogenic RAS and the p53 tumor suppressor. PLoS Biol. 2008;6:2853–2868. doi: 10.1371/journal.pbio.0060301

22. Krtolica A, Parrinello S, Lockett S, Desprez PY, Campisi J. Senescent fibroblasts promote epithelial cell growth and tumorigenesis: a link between cancer and aging. Proc Natl Acad Sci U S A. 2001;98:12072–12077. doi: 10.1073/pnas.211053698

23. Storer M, Mas A, Robert-Moreno A, Pecoraro M, Ortells MC, Di Giacomo V, Yosef R, Pilpel N, Krizhanovsky V, Sharpe J, et al. Senescence is a developmental mechanism that contributes to embryonic growth and patterning. Cell. 2013;155:1119–1130. doi: 10.1016/j.cell.2013.10.041

24. Yao H, Wallace J, Peterson AL, Scaffa A, Rizal S, Hegarty K, Maeda H, Chang JL, Oulhen N, Kreiling JA, et al. Timing and cell specificity of senescence drives postnatal lung development and injury. Nature Communications. 2023;14:273. doi: 10.1038/s41467-023-35985-4

25. Narni-Mancinelli E, Chaix J, Fenis A, Kerdiles YM, Yessaad N, Reynders A, Gregoire C, Luche H, Ugolini S, Tomasello E, et al. Fate mapping analysis of lymphoid cells expressing the NKp46 cell surface receptor. Proceedings of the National Academy of Sciences. 2011;108:18324–18329. doi: doi:10.1073/pnas.1112064108

26. Madisen L, Zwingman TA, Sunkin SM, Oh SW, Zariwala HA, Gu H, Ng LL, Palmiter RD, Hawrylycz MJ, Jones AR, et al. A robust and high-throughput Cre reporting and characterization system for the whole mouse brain. Nature Neuroscience. 2010;13:133–140. doi: 10.1038/nn.2467

27. Kliment CR, Englert JM, Crum LP, Oury TD. A novel method for accurate collagen and biochemical assessment of pulmonary tissue utilizing one animal. Int J Clin Exp Pathol. 2011;4:349–355.

28. Soutiere SE, Mitzner W. On defining total lung capacity in the mouse. Journal of Applied Physiology. 2004;96:1658–1664. doi: 10.1152/japplphysiol.01098.2003

29. Virshup I, Rybakov S, Theis FJ, Angerer P, Wolf FA. anndata: Access and store annotated data matrices. The Journal of Open Source Software. 2024;9:4371.

30. Wolf FA, Angerer P, Theis FJ. SCANPY: large-scale single-cell gene expression data analysis. Genome Biology. 2018;19:15. doi: 10.1186/s13059-017-1382-0

31. Young MD, Behjati S. SoupX removes ambient RNA contamination from droplet-based single-cell RNA sequencing data. GigaScience. 2020;9. doi: 10.1093/gigascience/giaa151

32. Germain PL, Lun A, Garcia Meixide C, Macnair W, Robinson MD. Doublet identification in single-cell sequencing data using scDblFinder. F1000Res. 2021;10:979. doi: 10.12688/f1000research.73600.2

33. Traag VA, Waltman L, van Eck NJ. From Louvain to Leiden: guaranteeing well-connected communities. Scientific Reports. 2019;9:5233. doi: 10.1038/s41598-019-41695-z

34. Korsunsky I, Millard N, Fan J, Slowikowski K, Zhang F, Wei K, Baglaenko Y, Brenner M, Loh P-r, Raychaudhuri S. Fast, sensitive and accurate integration of single-cell data with Harmony. Nature Methods. 2019;16:1289–1296. doi: 10.1038/s41592-019-0619-0

35. Finak G, McDavid A, Yajima M, Deng J, Gersuk V, Shalek AK, Slichter CK, Miller HW, McElrath MJ, Prlic M, et al. MAST: a flexible statistical framework for assessing transcriptional changes and characterizing heterogeneity in single-cell RNA sequencing data. Genome Biology. 2015;16:278. doi: 10.1186/s13059-015-0844-5

36. Butler A, Hoffman P, Smibert P, Papalexi E, Satija R. Integrating single-cell transcriptomic data across different conditions, technologies, and species. Nature Biotechnology. 2018;36:411–420. doi: 10.1038/nbt.4096

37. Hänzelmann S, Castelo R, Guinney J. GSVA: gene set variation analysis for microarray and RNA-Seq data. BMC Bioinformatics. 2013;14:7. doi: 10.1186/1471-2105-14-7

38. Ritchie ME, Phipson B, Wu D, Hu Y, Law CW, Shi W, Smyth GK. limma powers differential expression analyses for RNA-sequencing and microarray studies. Nucleic Acids Research. 2015;43:e47–e47. doi: 10.1093/nar/gkv007

39. Jin S, Plikus MV, Nie Q. CellChat for systematic analysis of cell-cell communication from single-cell transcriptomics. Nat Protoc. 2025;20:180–219. doi: 10.1038/s41596-024-01045-4

40. Phipson B, Sim CB, Porrello ER, Hewitt AW, Powell J, Oshlack A. propeller: testing for differences in cell type proportions in single cell data. Bioinformatics. 2022;38:4720–4726. doi: 10.1093/bioinformatics/btac582

41. Yu G, Wang LG, Han Y, He QY. clusterProfiler: an R package for comparing biological themes among gene clusters. Omics. 2012;16:284–287. doi: 10.1089/omi.2011.0118

42. Warburton D, El-Hashash A, Carraro G, Tiozzo C, Sala F, Rogers O, De Langhe S, Kemp PJ, Riccardi D, Torday J, et al. Lung organogenesis. Curr Top Dev Biol. 2010;90:73–158. doi: 10.1016/s0070-2153(10)90003-3

43. James J, Dekan A, Niihori M, McClain N, Varghese M, Bharti D, Lawal OS, Padilla-Rodrigez M, Yi D, Dai Z, et al. Novel Populations of Lung Capillary Endothelial Cells and Their Functional Significance. Res Sq. 2023. doi: 10.21203/rs.3.rs-2887159/v1

44. Hernandez-Segura A, Nehme J, Demaria M. Hallmarks of Cellular Senescence. Trends Cell Biol. 2018;28:436–453. doi: 10.1016/j.tcb.2018.02.001

45. Sharpless NE, Sherr CJ. Forging a signature of in vivo senescence. Nat Rev Cancer. 2015;15:397–408. doi: 10.1038/nrc3960

46. Binet F, Cagnone G, Crespo-Garcia S, Hata M, Neault M, Dejda A, Wilson AM, Buscarlet M, Mawambo GT, Howard JP, et al. Neutrophil extracellular traps target senescent vasculature for tissue remodeling in retinopathy. Science. 2020;369:eaay5356. doi: doi:10.1126/science.aay5356

47. Muñoz-Espín D, Cañamero M, Maraver A, Gómez-López G, Contreras J, Murillo-Cuesta S, Rodríguez-Baeza A, Varela-Nieto I, Ruberte J, Collado M, et al. Programmed Cell Senescence during Mammalian Embryonic Development. Cell. 2013;155:1104–1118. doi: 10.1016/j.cell.2013.10.019

48. Mavria G, Vercoulen Y, Yeo M, Paterson H, Karasarides M, Marais R, Bird D, Marshall CJ. ERK-MAPK signaling opposes Rho-kinase to promote endothelial cell survival and sprouting during angiogenesis. Cancer Cell. 2006;9:33–44. doi: 10.1016/j.ccr.2005.12.021

49. Zamai L, Ahmad M, Bennett IM, Azzoni L, Alnemri ES, Perussia B. Natural killer (NK) cell-mediated cytotoxicity: differential use of TRAIL and Fas ligand by immature and mature primary human NK cells. J Exp Med. 1998;188:2375–2380. doi: 10.1084/jem.188.12.2375

50. Wolf KG, Crawford EB, Wartan NM, Schneiderman SK, Riehl VE, Dambaeva SV, Beaman KD. Ephrin-B2-expressing natural killer cells induce angiogenesis. JVS Vasc Sci. 2022;3:336–344. doi: 10.1016/j.jvssci.2022.08.003

51. Manaster I, Gazit R, Goldman-Wohl D, Stern-Ginossar N, Mizrahi S, Yagel S, Mandelboim O. Notch activation enhances IFNγ secretion by human peripheral blood and decidual NK cells. Journal of Reproductive Immunology. 2010;84:1–7. doi: 10.1016/j.jri.2009.10.009

52. Kuhn A, Brachtendorf G, Kurth F, Sonntag M, Samulowitz U, Metze D, Vestweber D. Expression of Endomucin, a Novel Endothelial Sialomucin, in Normal and Diseased Human Skin. Journal of Investigative Dermatology. 2002;119:1388–1393. doi: 10.1046/j.1523-1747.2002.19647.x

53. Shirazi SP, Negretti NM, Jetter CS, Sharkey AL, Garg S, Kapp ME, Wilkins D, Fortier G, Mallapragada S, Banovich NE, et al. Bronchopulmonary Dysplasia with Pulmonary Hypertension Associates with Loss of Semaphorin Signaling and Functional Decrease in FOXF1 Expression. *bioRxiv*. 2024:2024.2008.2027.609955. doi: 10.1101/2024.08.27.609955

54. Liu B, Zhu X, Kong L, Wang M, Spanoudis C, Chaturvedi P, George V, Jiao J-a, You L, Egan JO, et al. Bifunctional TGF-β trap/IL-15 protein complex elicits potent NK cell and CD8+ T cell immunity against solid tumors. Molecular Therapy. 2021;29:2949–2962. doi: 10.1016/j.ymthe.2021.06.001

55. Wei XW, Zhang YC, Wu F, Tian FJ, Lin Y. The role of extravillous trophoblasts and uterine NK cells in vascular remodeling during pregnancy. Front Immunol. 2022;13:951482. doi: 10.3389/fimmu.2022.951482

56. Li M, Li C, Liu YH, Xing Y, Hu L, Borok Z, Kwong KY, Minoo P. Mesodermal deletion of transforming growth factor-beta receptor II disrupts lung epithelial morphogenesis: cross-talk between TGF-beta and Sonic hedgehog pathways. J Biol Chem. 2008;283:36257–36264. doi: 10.1074/jbc.M806786200

57. Chen H, Zhuang F, Liu Y-H, Xu B, del Moral P, Deng W, Chai Y, Kolb M, Gauldie J, Warburton D, et al. TGF-β receptor II in epithelia versus mesenchyme plays distinct roles in the developing lung. European Respiratory Journal. 2008;32:285–295. doi: 10.1183/09031936.00165407

58. Hammond JD, Kielt MJ, Conroy S, Lingappan K, Austin ED, Eldredge LC, Truog WE, Abman SH, Nelin LD, Guaman MC. Exploring the Association of Male Sex With Adverse Outcomes in Severe Bronchopulmonary Dysplasia: A Retrospective, Multicenter Cohort Study. CHEST. 2024;165:610–620. doi: 10.1016/j.chest.2023.10.020

59. Binet ME, Bujold E, Lefebvre F, Tremblay Y, Piedboeuf B. Role of gender in morbidity and mortality of extremely premature neonates. Am J Perinatol. 2012;29:159–166. doi: 10.1055/s-0031-1284225

60. Vrijlandt EJ, Gerritsen J, Boezen HM, Duiverman EJ. Gender differences in respiratory symptoms in 19-year-old adults born preterm. Respir Res. 2005;6:117. doi: 10.1186/1465-9921-6-117

61. Zhang L, Pitcher LE, Prahalad V, Niedernhofer LJ, Robbins PD. Targeting cellular senescence with senotherapeutics: senolytics and senomorphics. Febs j. 2023;290:1362–1383. doi: 10.1111/febs.16350

62. Wickramasinghe LC, van Wijngaarden P, Johnson C, Tsantikos E, Hibbs ML. An Experimental Model of Bronchopulmonary Dysplasia Features Long-Term Retinal and Pulmonary Defects but Not Sustained Lung Inflammation. Frontiers in Pediatrics. 2021;9. doi: 10.3389/fped.2021.689699

